# The ‘very moment’ when UDG recognizes a flipped-out uracil base in dsDNA

**DOI:** 10.1101/2024.09.13.612628

**Authors:** Vinnarasi Saravanan, Nessim Raouraoua, Guillaume Brysbaert, Stefano Giordano, Marc F. Lensink, Fabrizio Cleri, Ralf Blossey

## Abstract

Uracil-DNA glycosylase (UDG) is the first enzyme in the base-excision repair (BER) pathway, acting on uracil bases in DNA. How UDG finds its targets has not been conclusively resolved yet. Based on available structural and other experimental evidence, two possible pathways are under discussion. In one, the action of UDG on the DNA bases is believed to follow a ‘pinch-push-pull’ model, in which UDG generates the base-flip in an active manner. A second scenario is based on the exploitation of bases flipping out thermally from the DNA. Recent molecular dynamics (MD) studies of DNA in trinucleosome arrays have shown that base-flipping can be readily induced by the action of mechanical forces on DNA alone. This alternative mechanism could possibly enhance the probability for the second scnenario of UDG-uracil interaction via the formation of a recognition complex of UDG with flipped-out base. In this work we describe DNA structures with flipped-out uracil bases generated by MD simulations which we then subject to docking simulations with the UDG enzyme. Our results for the UDG-uracil recognition complex support the view that base-flipping induced by DNA mechanics can be a relevant mechanism of uracil base recognition by the UDG glycosylase in chromatin.

## Introduction

The recognition and the subsequent cleavage of wrongly incorporated or chemically induced uracil bases in DNA by repair enzymes is the initial step in the base-excision repair (BER) process. It is performed by the enzyme Uracil-DNA glycosylase (UDG), a protein of 313 amino acids (UniProt P13051 UNG_HUMAN) that was the first glycosylase to be structurally characterized in complex with damaged DNA^1^; for a review of structural and functional properties of the UDG superfamily of glycosylases, see^2^. UDG structures were repeatedly resolved experimentally in interaction with DNA and made available trough the Protein Data Bank (PDB, https://www.rcsb.org).

UDG initiates the BER pathway for uracil excision by hydrolyzing a glycosylic bond between a uracil base and the deoxyribose sugar in order to produce a free uracil and an abasic site in the DNA strand. Human UDG also removes uracil from single-stranded DNA, but it is not active against uracil in RNA^3^. Uracil excision requires the base to be flipped out of the DNA double strand so that it can be processed in the enzyme active site^1^.

DNA glycosylases must be extremely efficient to inspect all bases in the genome (3×10^9^ base pairs) and detect and repair damaged bases (around 1 in 10^6^ bases per cell daily) before the genetic code is permanently altered in the next cycle of DNA replication. Given that there are on the order of 10^5^ copies of glycosylase molecules in the nucleus, each such protein excluding redundancy should be able to sample around 70,000 base pairs of DNA in every cell cycle (about 12 to 24 hours).^4^ The fact that, in practice, glycosylases can achieve their job much faster points to the existence of better than purely stochastic mechanisms.

How UDG actually finds the wrongly incorporated base is a question that cannot be conclusively answered by crystallography, as it obviously is a highly dynamic process and X-ray only allows for access to final stable states of the interacting species. In order to cleave the base, UDG must first gain access to it. Several structural studies found that different DNA repair proteins can actively flip out their target base extra-helically (the so-called ‘enzymatic’ base flipping, see e.g.^5–7^). Recent single-molecule experiments have provided evidence that UDG works in a ‘scanning’ mode, i.e. processing along DNA, combined with a ‘peaking’-mode, which involves the insertion of an amino acid into the DNA stack^8^. Already before experimental evidence had been provided for hopping and sliding motions of UDG on DNA^9^, a detailed ‘pinch-push-pull’ scenario had been formulated, in which the non-specific scanning step of UDG along DNA is followed by highly specific actions of UDG on DNA^3^. In this scenario, UDG first induces a kink in the DNA backbone, allowing for the formation of specific amino acid contacts with the base, which then leads to a backbone compression that ‘pinches’ the DNA. Subsequently, the intercalation loop of UDG penetrates the minor groove and flips out the uracil base. In the final ‘pull’ step, the glycosidic bond is cleaved. Importantly, apart from the non-specific diffusion or hopping along DNA, the amino-acid-DNA interaction is actively pursued by UDG. This ‘pinch-push-pull’ scenario thus starts from an initially pristine DNA, and requires UDG to very actively search for the ‘hidden’ base. There is good reason to doubt that search processes are the only, or even the dominant mechanism in locating wrong bases for excision in DNA. Based on available data from in-vitro experiments, a simple Monte Carlo (MC) random search simulation could well accommodate the experimental situation; however, it remains unrealistic by several orders of magnitude for realistically estimated chromatin densities in the eukaryotic nucleus (see Supplementary Material). Such considerations therefore motivate the investigation for alternative mechanisms, by which defect-containing DNA fragments may become highly visible, and more easily identifable by the repair proteins compared to serendipitous random search.

It is still under debate whether the spontaneous base flipping, which normally occurs at random all along the DNA because of thermodynamical fluctuations^10–13^, could be an even stronger attractor for glycosylases and other similarly functioning enzymes.^14–16^ Given the much longer lifetimes measured by NMR for extrahelical flipping (in the *µ*s to ms range) compared to intrahelical flip, and the fact that even partially flipped bases may become accessible^17, 18^, spontaneous flipping must therefore be properly investigated as a possible first-step, or possibly the very rate-limiting event, in the glycosylase search for damaged DNA sites. Base flipping in pristine DNA in fact happens stochastically and an encounter with UDG could therefore also happen randomly; it is however easy to estimate that the probability of a purely random encounter (product of two independent random events), even with a substantial concentration of UDG molecules, will be generally too low to be of practical importance. DNA in the cell nucleus, however, is subject to numerous untargeted interactions, which might indeed lead to a large increase in the rate of base flipping, compared to the thermally-assisted one, thereby largely facilitating uracil recognition by UDG. In particular, mechanical constraints acting on the DNA structure have been indicated as an additional source of localized bending, twisting and kinking of the double helix, all which are favorable conditions for increasing the base-flipping rate. For example, in a recent work^19^, we modelled by fully-atomistic molecular dynamics (MD) simulations an array of three nucleosomes, under applied mechanical forces in a range reminding the compressive regime that cells may be typically subject during their lifetime. These simulations have shown that the resulting compression of the structure can induce kink instabilities in the linker DNA which facilitate the opening of the double-stranded linker DNA and the flipping-out of the bases^19^. This mechanical mechanism leads to the formation of localized denaturation bubbles through a mechanical twist of the DNA double helix^20^. It can therefore be concluded that the ubiquitous mechanical action on DNA could therefore present a partially or fully exposed uracil base to UDG. This observation thus raises the question whether DNA mechanics can play, among other effects, a relevant role in the recognition process of uracil bases by repair enzymes.

Starting from the insights gained in^19^, in this paper we characterize the interaction of UDG with exposed uracil bases and attempt to find the instant in which the recognition complex is formed between the flipped-out base and UDG. In other words, we aim at finding the ‘very moment’ (a notion inspired by^21^) when UDG can recognize a flipped-out base without having previously intervened in its formation. Here we study this problem by a combination of the controlled induction of uracil base-flipping via MD and protein-DNA docking simulations. Our finding of this ‘very moment’ of encounter entails the question of the quality of the complex formed: we compare our simulation results to the crystal structure of the UDG-DNA complex, which is assumed as representative for the action of the enzyme on uracil.

The flipping of nucleic bases has been studied by MD simulations by several authors, see, e.g.^22–25^. Studying the spontaneous base flipping is challenging, as the free energy penalty for the extrahelical state of individual DNA bases is about 10 kcal/mol^26, 27^. According to previous nuclear magnetic resonance (NMR) studies^10, 28^, the lifetime of the extrahelical state of a flipped DNA base is on the order of microseconds (µs). In contrast, the intrahelical state lasts from milliseconds to hundreds of milliseconds, depending entirely on the stability of distinct base pairs in the dsDNA. This significant difference between the timescales accessible in all-atom molecular dynamics (MD) simulations and those of base flipping makes it difficult to obtain converged statistics in computer simulations. Therefore, the rare-event computational methodology is required for spontaneous base flipping mechanism. Várnai and Lavery^22^, Huang et al.^29^ and Law and Feig^30^ have used external forces to induce individual base flipping through umbrella sampling and replica exchange simulation methodologies. In this work, we have used collective variable (CV) based metadynamics^31^ simulations to study the base-flipping mechanism.

Having the base-flipped DNA structures at hand, either from the dedicated simulations described here, or from our earlier work on the compression of trinucleosomes^19^, we use protein-DNA docking methods to study the formation of the recognition complexes as a function of the flipping of the base. Contrary to protein-protein docking, where standard methods working in most of the cases are established, protein-DNA docking requires special attention to the algorithm used, in particular when DNA bears uracil. It is even more essential to assess the efficiency of these different methods for our system as it corresponds to a non-standard case: the nature of the DNA strands that are docked to UDG here is highly altered, as the treatment done to induce base flipping is causing deformations on the strand, and the flipped-out base represents a significant conformational change of the pristine dsDNA. In order to determine the best-suited protein-DNA docking method for our case, we had to test several available algorithms, and describe in detail our quantification of the quality of the encounter complex through protein-DNA docking.

## Results

### Mechanisms of uracil base flipping and associated structural deformations

We have obtained dsDNA conformations with flipped uracil bases in two ways. The first approach makes use of the structures generated during the compression MD simulations of tricnucleosome arrays performed in our earlier work^19^. How these structures were selected and prepared for our docking simulations is discussed in Methods.

In a second approach, we have generated flipped-out bases in a controlled way from mutated dsDNA. For this we used a dsDNA 17-mer in the standard B-DNA form given by the sequence 5’–CAGGATGTATATATCTG–3’. The thymine nucleotide T at position 12 was mutated to a uracil base with its cartesian coordinates generated by the Web 3DNA server^32^. The targeted central base pair for flipping was U_12_:A_23_, with U_12_ as the target nucleotide for base flipping; the geometry is illustrated in Figure 1 a). First, an all-atom conventional MD simulation was performed to stabilize the dsDNA system over approximately 100 ns. The root-mean-square deviation (RMSD) was used to monitor the overall stability of the dsDNA, as shown in Figure 1 b). The overall RMSD fluctuation was around 1-3 Å, which shows that the relative structural changes are very small. Moreover, the root mean-square fluctuation (RMSF) values calculated for the uracil-mutated dsDNA reveal very small fluctuations at the individual base level, see Figure 1 c).

**Figure 1.**
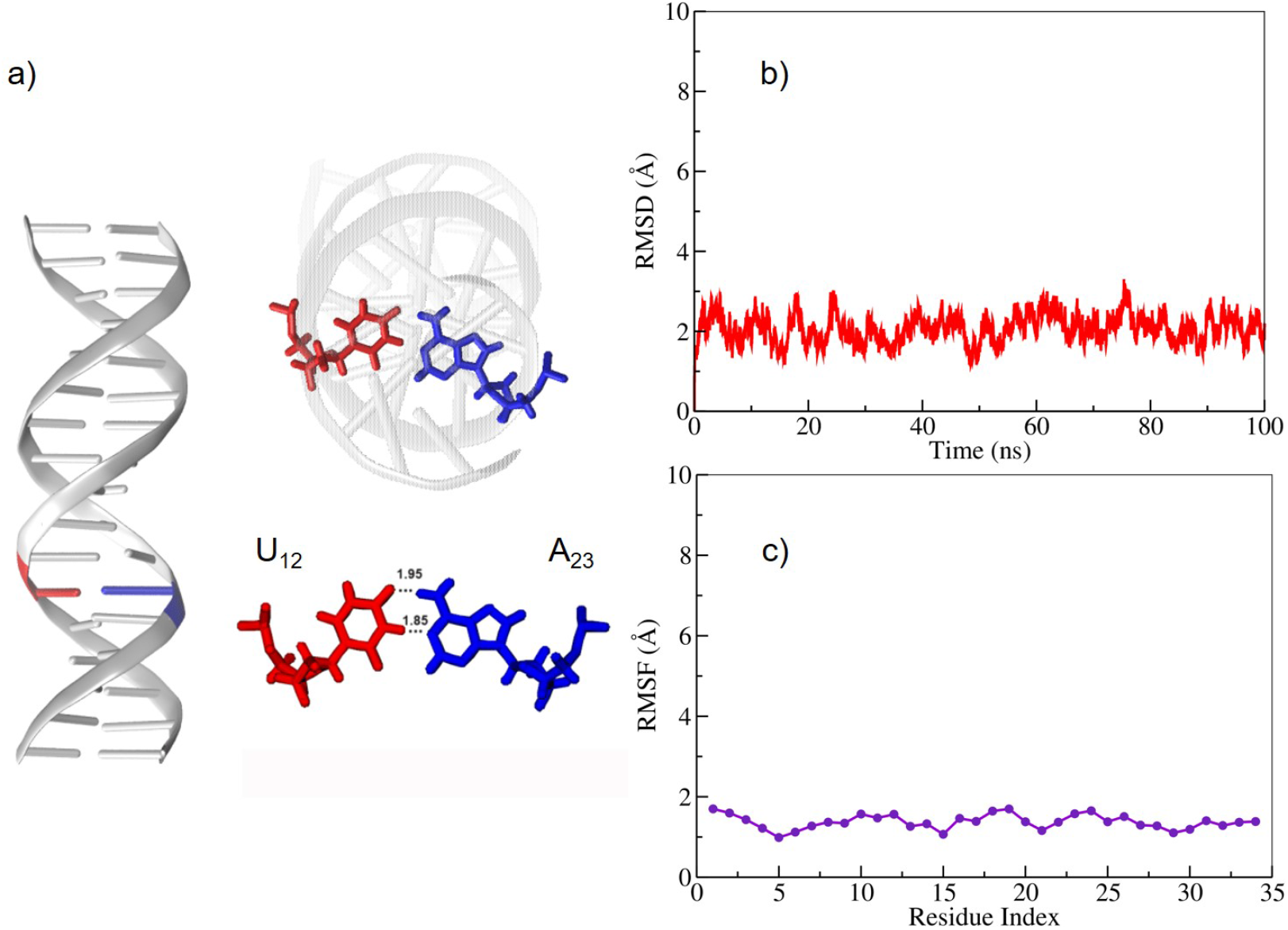
a) Uracil-mutated dsDNA system with red and blue colors representing uracil and adenine residues, respectively. b) Root mean square deviation (RMSD) and c) Root mean square fluctutation (RMSF) of the uracil-mutated dsDNA system during the first 100 ns of the MD trajectory.

During the base flipping simulation, an external history-dependent bias potential is applied to the system which can be expressed as a sum of Gaussians along the collective variables (CV) to enhance the sampling efficiency. The CVs for the uracil base flip have been chosen following existing literature, *i*.*e*. different versions of the center-of-mass pseudo-dihedral angle (CPD), see^33^ and CPDa/b^24^. For further details, see Methods.

The metadynamics simulations were conducted for the uracil mutated dsDNA system for about 100 ns. The pseudo-dihedral angles (CPD and CPDb) were used as the coordinates for free energy mapping to describe the uracil flipped-in and flipped-out states using the potential of mean force (PMF) analysis as shown in Figure 2 a). The base opening pathway snapshots are shown in Figure 2 b). In this Figure, notable similarities and differences between the two energy profile schemes are observed. The global minima of uracil embedded states at approximately 10° in CPDb correlate well with previous theoretical studies^34^, whereas in the CPD scheme, it is approximately 40°. Here, the preselected CPD reaction coordinates scheme yields slightly different results. Both PMF profile schemes display a separation into two distinct regions: the major groove and the minor groove. A nucleobase within the dsDNA aligns with CPDb > 0° when flipping into the major groove pathway, whereas it aligns with CPDb < 0° when flipping into the minor groove pathway. In Figure 2 a), the spontaneous flipping of uracil through the major groove has a lower free energy barrier than through the minor groove in both CPDb and CPD schemes, respectively. The lower free energy barriers and their corresponding base opening angles were about 7.5 kcal/mol and 30° for CPDb and 6 kcal/mol and 60° for CPD schemes, respectively. This observation is in good agreement with the previous meta-eABF simulation^34^. The flipped-out pseudo dihedral angle uracil was about 180°, and the corresponding free energy barriers were about 8.2 kcal/mol and 9 kcal/mol for CPDb and CPD schemes, respectively. The transition from the flipped-in to the flipped-out state of uracil has a higher free energy barrier in the CPD than in the CPDb schemes. The obtained results suggest that the observed differences could be attributed to the effect of the reaction coordinate definition on the details of the PMF. Furthermore, the results indicate that uracil flips out more frequently through the major groove pathway than through the minor groove pathway.

**Figure 2.**
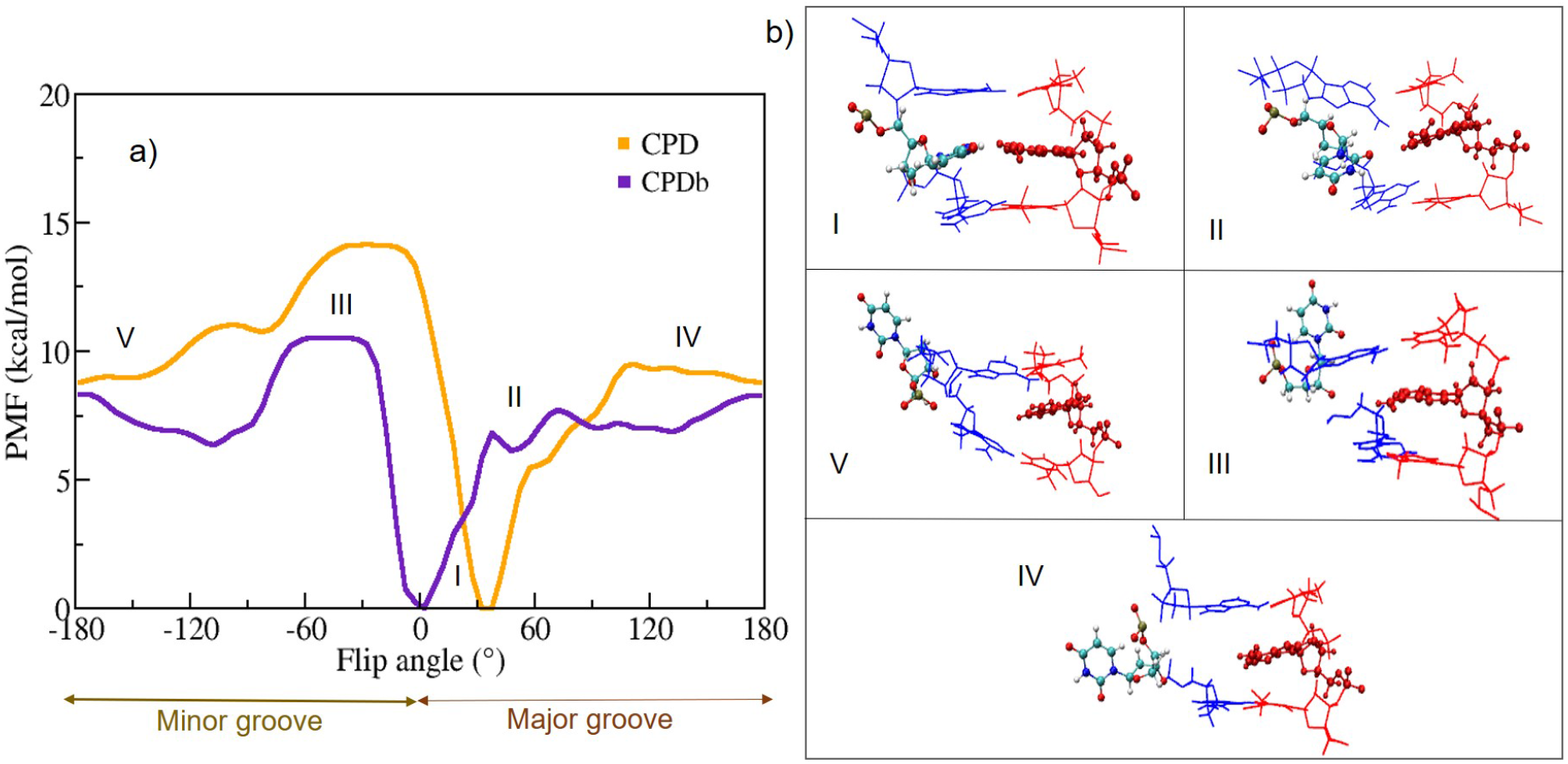
a) Free energy profile (Potential of Mean Force, PMF) as a function of the Center Of Mass (COM) of the pseudo-dihedral angle for the uracil-mutated dsDNA; b) The representative snapshots for the uracil base flipping steps [here region I is the Watson-Crick base pair, II, III, V regions are the intermediate steps, and IV is the fully flipped state of uracil].

In order to understand the structural properties of uracil base opening, analyses of the hydrogen bond distances (N1⋯H3 and O4⋯H6’) and the center of mass (COM) of distance between uracil and adenine were performed for the metadynamics simulation trajectories, as displayed in Figure 3. At the beginning of the simulation, the COM distance between uracil and adenine bases were about 10.5 Å. During the simulation between 30-40 ns, this distance increases slightly to 12.5 Å. After 75 ns (Figure 3 c), the uracil base is completely flipped-out, causing the distance to increase to 17 Å, whereas the hydrogen bond (H-bond) distance between N1(A)⋯H3(U) and O4(U)⋯H6’(A) increased from 1.82 Å to 15.9 Å and from 2.12 Å to 18.4 Å, respectively; see Figure 3 a). In the CPDb (Figure 3 b) scheme, the COM distance increases from 10.5 to 15 Å, whereas the distances of the H-bonds N1⋯H3 and O4⋯H6’ have increased from 1.82 Å to 16.2 Å and from 2.08 Å to 16.4 Å, respectively.

**Figure 3.**
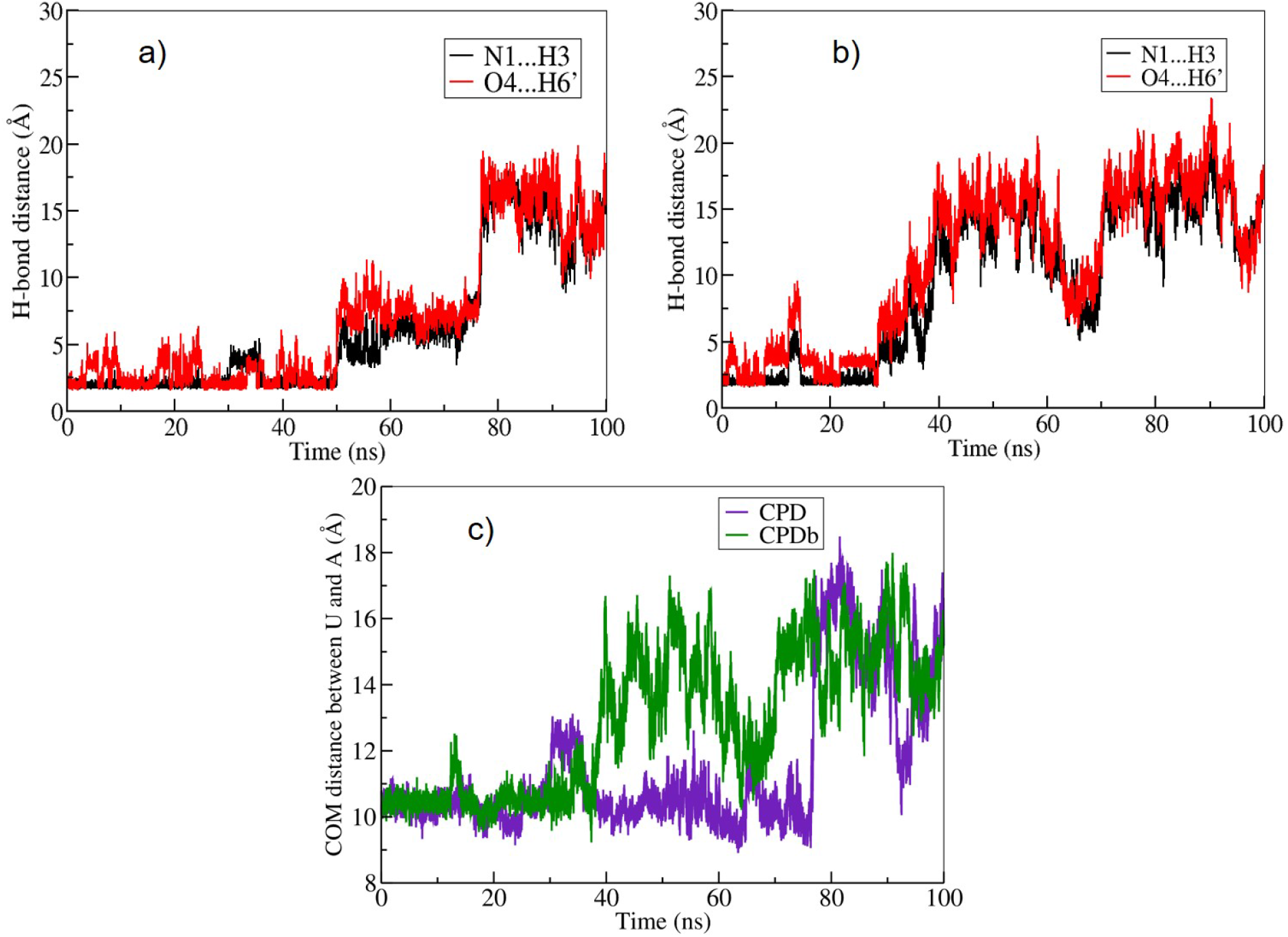
a) and b) the hydrogen bond distances between N1⋯H3 and O4⋯H6’ for the CPD and CPDb schemes during the course of the simulation, respectively; c) Likewise, the calculated average centre of mass of the distance between the uracil and adenine residues for CPD and CPDb schemes.

Further, we have analysed the uracil-mutated dsDNA structural deformations. For this we used the Curves+ program^35^. The results are given in Figure 4 and the Supplementary Figure S2. The intrabase parameters (buckle, opening) and the interbase parameter (tilt) for the flipped-in to flipped-out transition of uracil in the U.A base pair ranged from −2.5° to −57.4°, 10.7° to −168.2°, and −1.6° to 16.0°, respectively. The total dsDNA bending angle increased from 3.6° to 68.6°. In particular, the major and minor groove values of flipped-out uracil dsDNA were about 11 Å and 7 Å, respectively. Since a wider major groove was confirmed more favourable for base flipping, the observed increase in the major groove width favors a structural adaptation that facilitates uracil flipping in the DNA sequence^30^.

**Figure 4.**
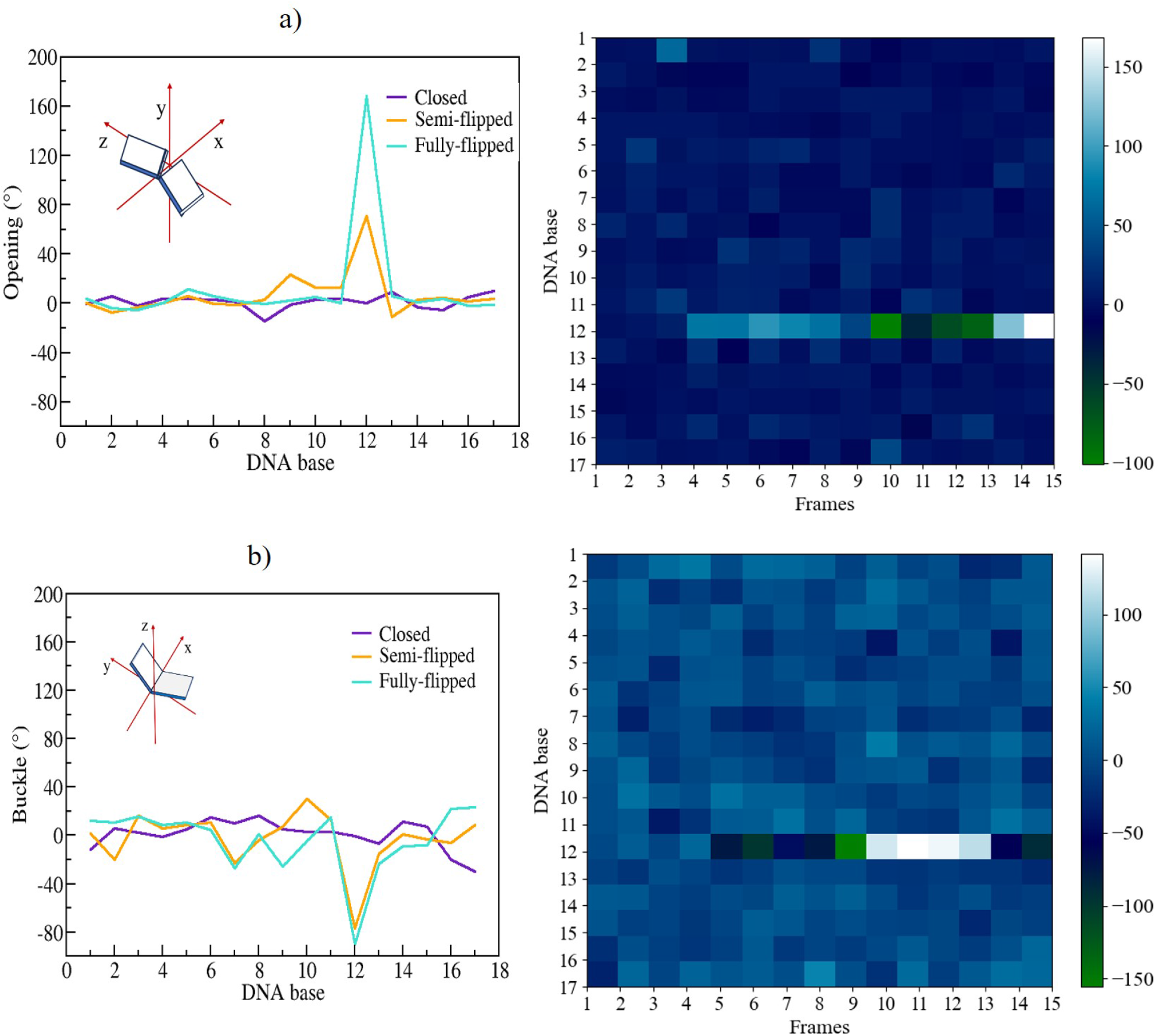
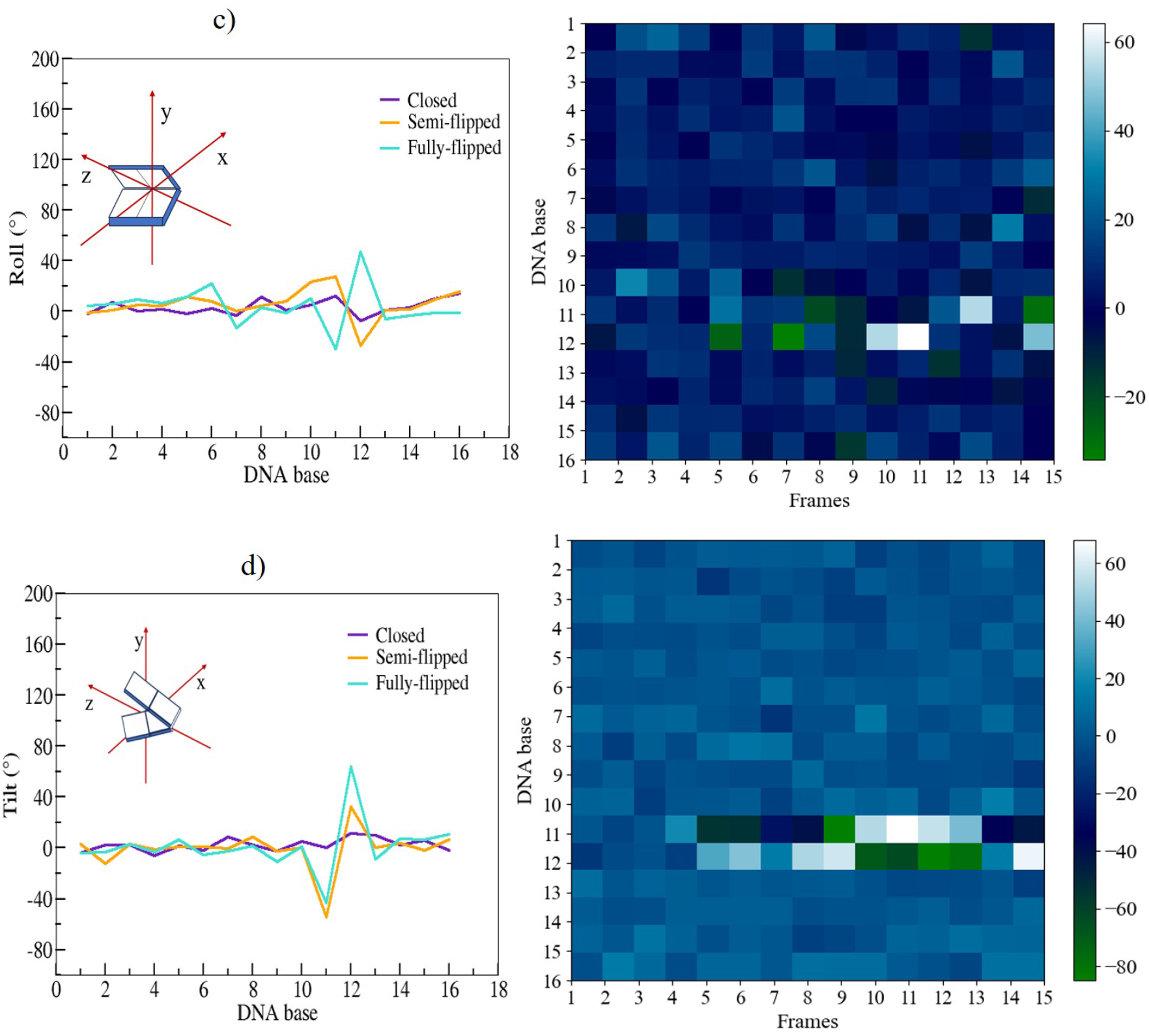
Intra-base pair parameters: a) Opening and b) Buckle, and inter-base pair parameters c) Roll and d) Tilt of uracil-mutated dsDNA for the CPDb scheme were calculated using the CURVES+ program. The plots on the right hand side are the respective heat maps of Opening, Buckle, Roll, and Tilt with respect to the trajectory frames. Here, the dark shades (blue to green) indicate negative values, and the light shades (cyan to white) indicate positive values of these quantities for the uracil-mutated dsDNA.

### Docking UDG to flipped uracil bases in dsDNA

Two isoforms of human UDG are known to originate from the alternative splicing of the UNG gene. The canonical isoform is UNG2 (Uniprot P13051-1) and has a length of 313 aa. It is expressed in the mitochondrion and is the one referred in the introduction as the first glycosylase characterized in complex with damaged DNA. The second isoform is UNG1 (Uniprot P13051-2, 304 aa), which is expressed in the nucleus and hence the isoform we used for our docking simulations. The difference between these two isoforms lies in their N-terminal beginning portions: UNG1 from residue 36 to its C-terminal end (304) is identical to UNG2 from residue 45 to its C-terminal end (residue 313). The structure of the predicted UNG1 structure we use is shown in Figure 5 a).

**Figure 5.**
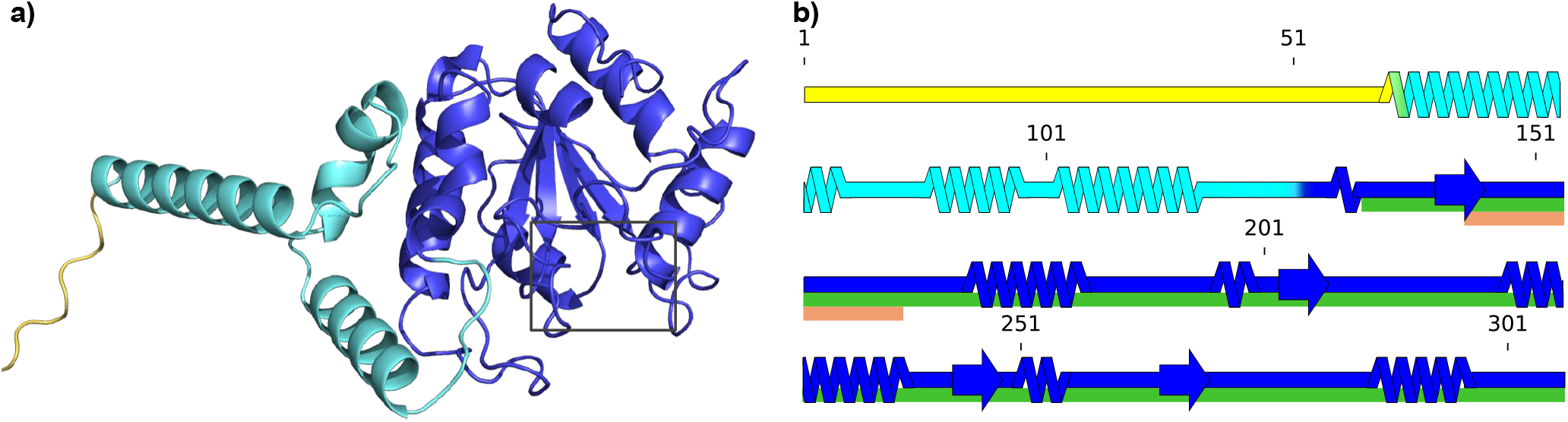
a) Cartoon representation of UDG tertiary structure predicted with Alphafold2 (AF2) as described in Methods. The structure is represented with PyMOL^36^. Its residues are colored according to the backbone qualitative flexibility evaluation described in the main text and in Methods. The three flexibility levels are: <u>yellow</u> for flexible, <u>blue</u> for rigid and <u>cyan</u> for intermediate level between flexible and rigid. In a), the black frame at the bottom right designates UDG’s catalytic pocket; b) 2D representation of UDG secondary structure with the same color code as in a), drawn with SSDraw^37^. Notable regions are highlighted under the concerned residues: the catalytic site is shown in <u>orange</u> (InterPro IPR018085) and the conserved UDG domain in <u>green</u> (InterPro IPR005122).

As a first step in studying the docking process we have investigated the flexibility of UDG, since the docking of proteins to DNA is clearly influenced by the flexibility of both docking partners. The results of our investigation are reported in Figure 5 b). The flexibility information obtained with the MEDUSA webserver^38^(see Methods) reveals that the portion containing UDG’s catalytic site is predicted as mostly rigid. Upstream of this rigid section the protein is less rigid, even ending on a disordered and highly flexible N-terminal tail.

To study protein-DNA interactions, several docking algorithms and programs have been developed and are mostly available either from a webserver or with its code made publicly available, see Methods.^39–44^. We have tested these docking programs on the crystal structure PDB:1EMH which we take as our reference structure for the recognition complex. The results of these docking attempts are summarized in Supplementary Figure S4. We found that pyDockDNA performed best because it manages to discriminate between uracil and thymine, and was therefore selected for our docking simulations (see Methods).

In order to have a global view of UDG’s encounter with the damaged DNA strand, we took snapshots from the MD simulations of either the trinucleosome structure, as obtained in our previous work^19^, or from the MD base-flipping simulations described above. We denote these structures by either a ‘T’ for trinucleosome or ‘O’ for the oligomer (17-mer) dsDNA in the following. Table 1 collects the results of the measurement of suitable angles and distances to characterize the base-flip structures selected for the docking process. We characterized the docked structure by a distance and an angle measurement which is described in Methods.

**Table 1.** Quantitative description of impacting features for the structures ‘T’ from the trinucleosome simulations and ‘O’ from the 17-mer dsDNA oligomers, resulting from a gradual base flipping with their docking success evaluation. The distance *d*_*U*−*groove*_ measurements and *Dihedral angle* metrics are described in Supplementary Figure S3. The angle evaluates the extent of the base flipping, the distance indicates the minimal distance between the base and both edges of the groove. The closer the base is from the backbone of the opposite strand, the more sterically hindered it is. The sign “−” in dihedral angles implicates flipping towards the minor-groove.

|  |  |  |  |  |  |  |  |  |
| --- | --- | --- | --- | --- | --- | --- | --- | --- |
|  | 1EMH |  |  |  |  |  |  |  |
| Dihedral angle (°) | −171.75 |  |  |  |  |  |  |  |
| $d_{U-groove}$ (Å) | 20.8 | | | | | | | |
| Success | **High** |  |  |  |  |  |  |  |
|  | T1 | T2 | T3 | T4 | T5 | T6 | T7 | T8 |
| Dihedral angle (°) | −5.85 | 2.61 | −4.71 | −3.16 | −8.66 | −66.12 | −96.58 | −83.18 |
| $d_{U-groove}$ (Å) | 11.8 | 12.9 | 14.2 | 11.5 | 12.9 | 12.0 | 10.4 | 10.8 |
| Success | None | None | None | None | None | Low | Medium | Medium |
|  | T9 | T10 | T11 | T12 |  |  |  |  |
| Dihedral angle (°) | −108.82 | 145.12 | −119.20 | 175.9 |  |  |  |  |
| $d_{U-groove}$ (Å) | 11.1 | 8.3 | 12.8 | 10.4 | | | | |
| Success | Medium | Medium | Medium | Medium |  |  |  |  |
|  | O1 | O2 | O3 | O4 | O5 | O6 | O7 | O8 |
| Dihedral angle (°) | 41.8 | 29.8 | 151.9 | −173.2 | −103.6 | 151.7 | 103.9 | 160.5 |
| $d_{U-groove}$ (Å) | 14.4 | 15.2 | 13.2 | 16.4 | 10.9 | 15.8 | 10.0 | 13.3 |
| Success | None | None | **High** | **High** | Medium | **High** | Low | **High** |

A common feature appearing for every successful docking, as shown in Table 1, is a sufficiently wide opening angle of the flipped-out uracil base and a reasonable distance measure (the lesser being 13 Å) to the opposite DNA strand’s backbone. Both conditions are to be fulfilled for UDG to have enough space to reach the damaged base. These two conditions appear to be the main factors for a high success in docking as compared to the less successful conformations.

To analyse the docking results, we first filtered the conformations sampled by pyDockDNA. From the initial 10,000 docked structures that pyDockDNA produces with FTDOCK, it selects the 100 best according to a scoring function, which in this case is the one with no desolvation. According to the Supplementary Figure S5 a), this associated score is not sufficient to differentiate between the good and the bad docking solutions.

In order to sort the predictions by relevance, i.e. by how well the obtained interface corresponds to the one observed in 1EMH, we devised a protocol described in Methods which measures the similarity of the uracil position between the docking structure and 1EMH used as our reference structure. This measure, represented in Figure 6 is then translated to an indicator listed in Table 1 as ‘success’. According to Figure 6, the DNA strand with the highest success with a near perfect similarity to 1EMH is ‘O4’. Its docking success coincide with the structure having the highest value in both the opening angle and the groove width according to Table 1.

**Figure 6.**
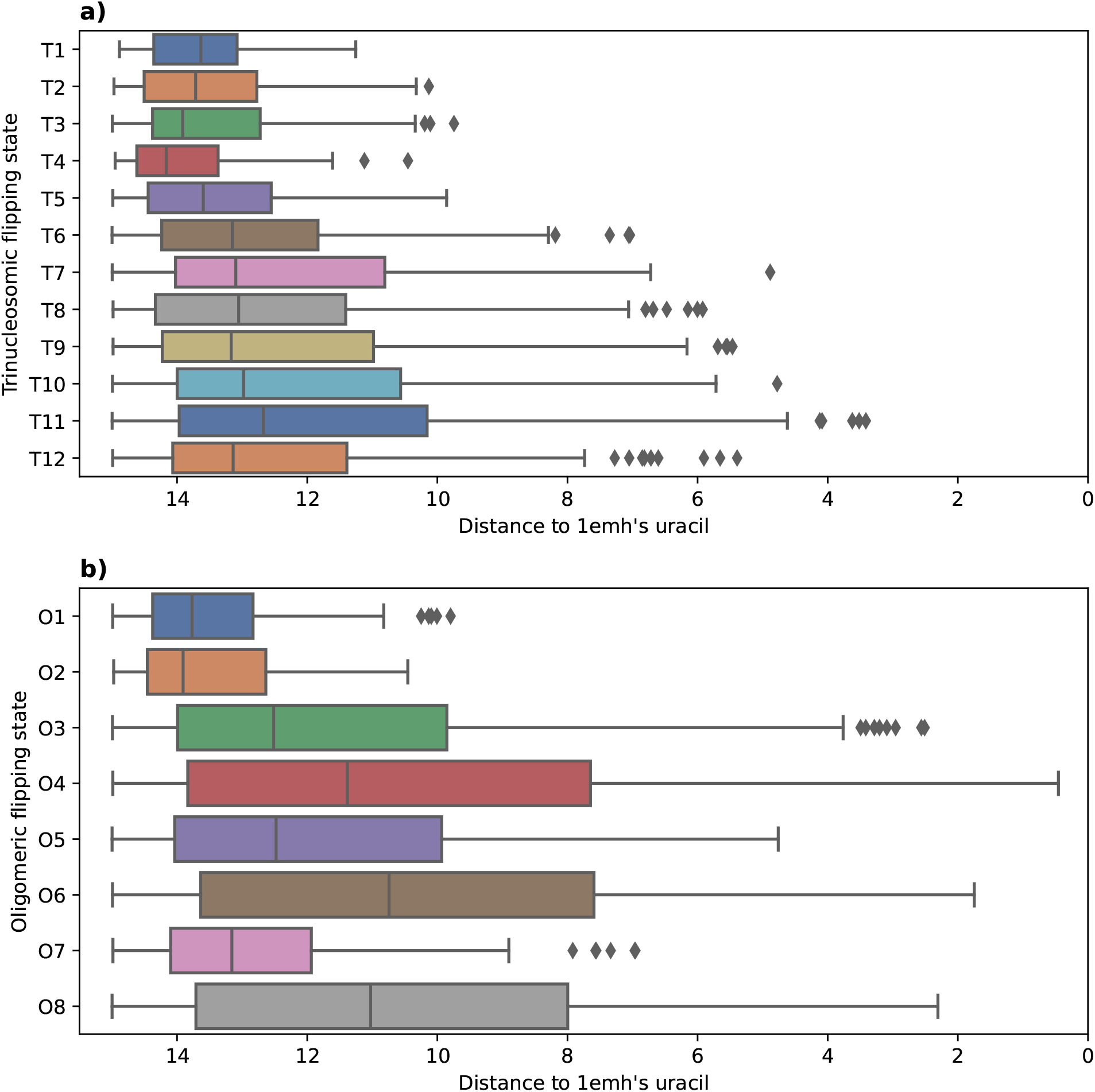
Distance of the docking-predicted uracil relative to the one of 1EMH, depending on the DNA structure used for docking with trinucleosomic strands in a) and oligomeric strands in b). The distance (Å) is computed after superimposing both UDG from the docked structure and from 1EMH. Only distances under 15 Å are conserved. X-axis goes from 15 Å to 0 Å. Distances are computed as described in Methods and illustrated in Supplementary Figure S6.

According to these measures, the highest docking success of both ‘T’ dsDNA and ‘O’ dsDNA (see Figure 6), respectively ‘T11’ and ‘O4’, were selected for illustration purposes in Figure 7. It denotes the high similarity of these docking successes to the 1EMH structure. ‘T12’ has a way wider flipping angle while being less successful than ‘T11’ (Figure 6 a)) for docking to UDG. The determining factor making ‘T11’ the most prone among the ‘T’ dsDNA to interacting with UDG seems to be the large distance between its uracil and the nearest groove’s end. On the other hand, ‘O4’ has at the same time the widest angle of base flipping and the most space in the groove making it the top pick for a successful docking.

**Figure 7.**
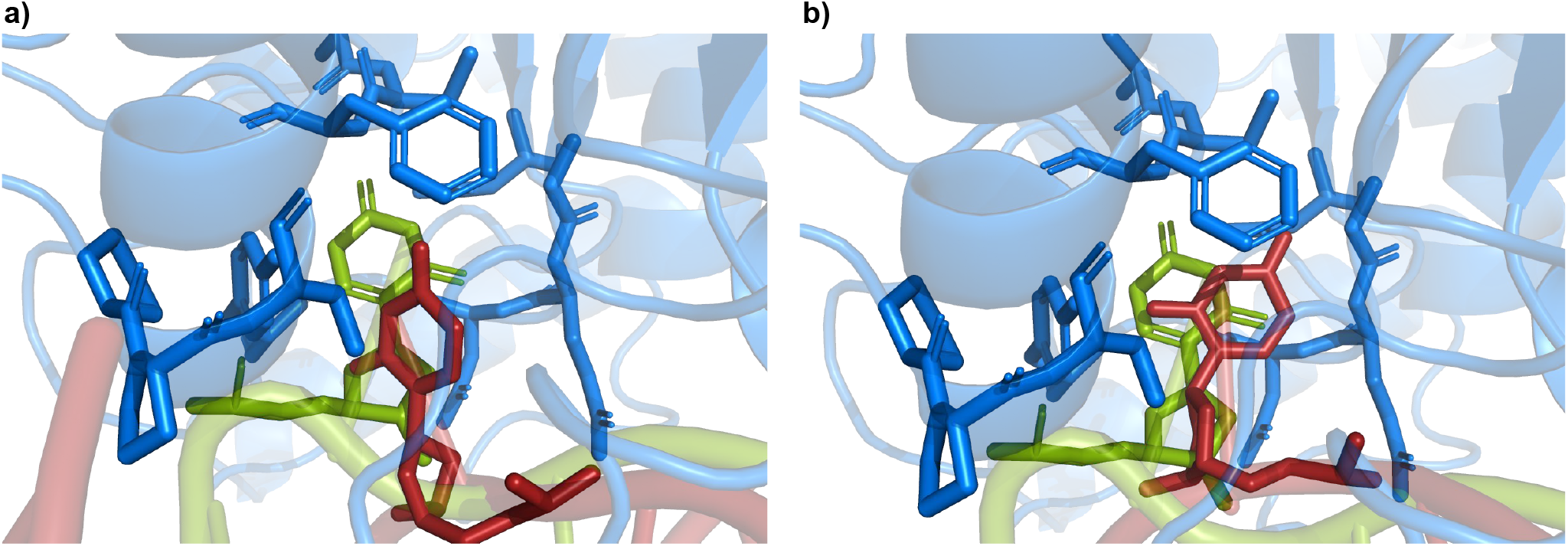
Docked structure for the highest success of both types of induced base flipping (‘T’ and ‘O’), identified according to Figure 6. a) ‘T11’ DNA strand (red) docked to UDG (blue; cartoon) with a focus on its catalytic pocket (blue; sticks). The uracil (red; sticks) from the docked DNA is compared to the one from 1EMH (green; sticks) as the reference. Following the same color code, b) shows the ‘O4’ DNA strand docked to UDG with 1EMH uracil as a reference to assess how well docking predicts the entrance of uracil into the catalytic site of UDG.

### Structural dynamics of uracil-flipped dsDNA bound to UDG enzyme

The optimal docking conformation of uracil-flipped dsDNA with the UDG enzyme (structure ‘O4’) was further explored using all-atom molecular dynamics simulations for 500 ns. The overall structural stability, compactness, and the flexibility of the complex system were evaluated using RMSD, radius of gyration (*R*_*g*_), and RMSF analyses. The mean RMSD values of the protein, the protein within the complex, and the protein-dsDNA complex were computed as 2.77 Å, 2.35 Å, and 3.14 Å, respectively, as shown in the Figure 8 a). The compactness of all three systems was assessed using *R*_*g*_ and the probability distribution of *R*_*g*_, as illustrated in the Figures 8 b) and 8 c). UDG initially exhibited higher fluctuations, around 20 Å, between 0 and 100 ns. After 100 ns, these fluctuations began to decrease to approximately 19.5 Å. However, after 350 ns, the UDG experienced increased fluctuations, primarily due to the loop present in the catalytic site (residues 210-220), which became more flexible and fluctuated significantly during the simulation. Consequently, the probability distribution of *R*_*g*_ adopted a bimodal conformation. In contrast, the *R*_*g*_ values for the UDG within the complex and the UDG-dsDNA system remained stable throughout the simulation, with values of 19.4 Å and 21 Å, respectively. The flexibility of the systems was further analysed using RMSF, as shown in the Figure 8 d). The flexibility of the loop present in UDG is higher than that in the complex system, as shown in the Figure 8 d). Notably, the loop residues (residues 145-158) located at the catalytic sites exhibit slight flexibility, with an RMSF of approximately 2.4 Å. The presence of uracil at the catalytic sites in the complex stabilizes the loop, rendering it more rigid due to the formation of hydrogen bonds with GLN144 (URA(O4)⋯H), ASP145 (URA(H5)⋯O), and PRO167 (URA(O2)⋯H) which are shown in Supplementary Figure S7. The loop residues 210-220 in the protein system are highly flexible, with an RMSF range of approximately 3.0-3.8 Å. These residues are further stabilized by the dsDNA during complex formation. In the complex, amino acids 272-278 exhibit higher fluctuations because the LEU272 residue in the loop intercalates into the nucleic acid strand to fill the gap created by uracil flipping into the catalytic site of the UDG enzyme, which correlates well with previous experimental findings^2^.

**Figure 8.**
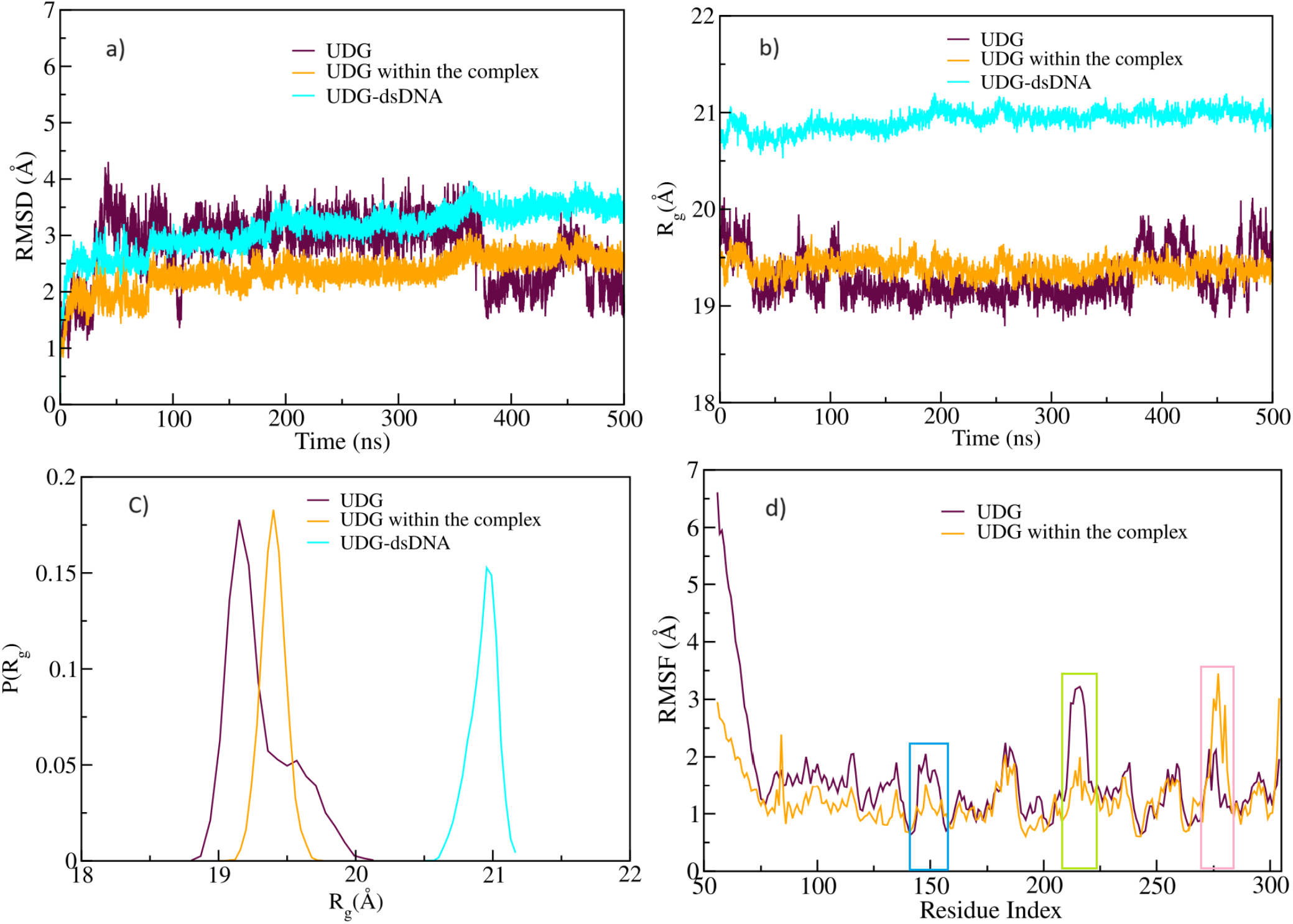
Structural analyses for the UDG-dsDNA complex. a) Root mean square deviation (RMSD), b) radius of gyration (*R*_*g*_), c) probability distribution of *R*_*g*_, and d) root mean square fluctuation (RMSF). Here, the blue box highlights the catalytic site of the UDG enzyme, the green box indicates the fluctuation of loop residues (residues 210 to 220), and the pink box indicates the fluctuation of LEU272 intercalated into the dsDNA within the UDG-dsDNA complex.

## Discussion

In this work we present arguments that the mechanism by which UDG identifies uracil bases in dsDNA cannot rely alone on a random search combined with the well-established ‘pinch-push-pull’-model that allows UDG to identify the presence of uracil inside the dsDNA strand. Uracil bases can present themselves to UDG via stochastic, thermally-induced base-flipping, but even more readily due to the ubiquitous presence of forces acting on dsDNA. As we have shown earlier^19^, this leads to instabilities of the dsDNA in the course of which bases flip out and become accessible to UDG. Here, we have studied whether this scenario is in accord with the base-flipping process of uracil from dsDNA, and whether the resulting structures can be properly recognized by UDG, i.e. whether a recognition complex between uracil and UDG can be formed. Our approach consisted in studying the base-flipping of uracil from dsDNA oligomers, using a collective variable/metadynamics approach within molecular dynamics simulations. In these simulations, uracil-flipped states were generated, passing through different stages of the flipping process in time. In parallel, we have selected base-flipped structure from our earlier large-scale MD simulations of trinucleosome arrays, in which base-flipping has occurred in the linker DNA of the arrays in the course of a mechanical instability in the compressed dsDNA. The opening angles of all of these structures have been quantified.

Subsequently, we have performed a study of rigid docking to the base-flipped uracil in dsDNA. While several protein-DNA docking programs have been developed in recent years, the study of protein-DNA docking is generally much less developed than the methodologies for protein-protein docking. In addition, the case of base-flipping requires the docking to be performed not towards a pristine, but towards a dynamically perturbed structure. This requirement has both necessitated a detailed study of available protein-dsDNA software as well as a careful investigation of the docking success, i.e. the formation of a ‘proper’ recognition complex. We found pyDockDNA to be the best suited software for our task. For the evaluation of the quality of the docking for our predicted recogition complexes, we took the crystal structure of UDG-dsDNA from PDB:1EMH as our reference structure. Our careful evaluation of the docking quality for both the oligomeric and trinucleosomic dsDNA is documented in Supplementary Figure S4. Overall, we identify two crucial factors in obtaining UDG-dsDNA recognition complexes, both of which are related to the opening angle of the uracil base. Firstly, we find that only a nearly fully flipped base (meaning between an angle of 150 and 180) allows for recognition complex formation. The second crucial factor is the width of the groove into which the base opens. The latter fact is well-known from earlier experimental studies of uracil embedded in nucleosomal DNA^45^. If these conditions are met we find that the formation of a high-quality recognition complex according to our criteria, as given in Methods, is possible. This is supported by MD simulations we performed on the best obtained complex.

Therefore, our study provides evidence that, besides the binding of UDG to dsDNA and its participation in a ‘pinch-push-pull’ mechanism, mechanical instabilities of dsDNA in the crowded nuclear environment that favor a destabilization of the dsDNA and therefore enhance the probability of base flipping beyond a merely random, thermally assisted flipping, may be of relevance as a pathway allowing the identification of uracil by UDG and thereby initializing the base-excision process. The initial BER recognition process is in fact occurring in a highly dynamic environment and very closely linked to DNA dynamics, at least in chromatin linker DNA. In future work we intend to look at the role of DNA dynamics in nucleosomes on the BER process.

## Methods

### Selecting and preparing the MD-snapshots ‘T’ from the trinucleosome simulations

We have selected twelve snapshots from the trinucleosome simulations described in^19^. They were chosen from the structure called T183, which was reconstructed by assembling three copies of the 197 nucleosome (PDB file 5NL0)^46^. More details as well as access to the trinucleosome structure can be found in^19^. The sequence of the dsDNA sequence extracted from that structure is given by 5’ −TACGUATGG −3’. In the original MD-simulations a fully paired dsDNA structure was used. The mechanically flipped-out base was replaced by uracil using the ‘Mutation’ section in the w3DNA 2.0 webserver.^47^

### MD simulations of uracil flipping in the dsDNA oligomer ‘O’

All atom molecular dynamics (MD) simulations were executed with the NAMD 2.14 package^48^ with the CHARMM 36 force fields^49^. A concentration of 0.15 mol/L Na^+^ and Cl^−^ ions were added at random positions to maintain the charge neutrality of the dsDNA system. The all-atom simulations employed periodic boundary conditions (PBC) and multiple time-stepping wherein local interactions were considered every 2 fs and full electrostatic evaluations were conducted every 2 time steps. The particle mesh Ewald method (PME) was employed for long-range electrostatic calculations^50^. The simulation box size was 79.7 × 40.6 × 40.9 Å^3^ with a distance of 10 Å between the box edges and the dsDNA. The cutoff and switching distances were set at 12 Å and 10 Å, respectively. Covalent bonds involving hydrogen were held rigid by the RATTLE^51^ and SETTLE algorithms^52^. The dsDNA system was minimized for 5000 steps using the conjugate gradient method. The pressure (constant NPT) was maintained at 1 atm, and the temperature was kept at 310 K. Temperature control was attained through Langevin dynamics for all non-hydrogen atoms, while pressure was controlled using a Nose-Hoover Langevin piston. The trajectories of the simulations were visualized and analysed by the visual molecular dynamic program (VMD 1.9.4)^53^ and the dsDNA conformational analysis were calculated by Curves+ program^35^.

### Choice of collective coordinates

In this study, the reaction coordinates have been described by three choices i) the centre-of-mass (COM) of pseudo-dihedral angle between the flipping base, the sugar group of the same nucleotide (group 1), the sugar group of the next nucleotide (group 2), the base of the next nucleotide plus the base of the opposing nucleotide (group 3), and the target base (group 4); ii) the center of the two flanking base pairs (group1), group 2 and 3 are defined by the flanking sugar moieties, and group 4 is defined by the target base for the flipping process; and iii) in the CPDb scheme, the group 1 is same as in CPDa, group 2 and 3 are defined by the flanking phosphate group, and the group 4 is defined by the target base for the flipping process. Among these choices, the uracil base opening event did not ensue with the CPDa scheme. As a result, all calculations described here were conducted using the CPD and CPDb schemes. The potential mean force was calculated for reaction coordinate between -180° and 180° into windows of 5° degree width; settings for Gaussian hillWeight = 0.001 kcal/mol and hillWidth = 2 bin width were added in the metadynamics simulation.

### Definition of docking partners

Docking here consists in simulating the interaction between UDG and the base-flipped DNA. The DNA strands used in these simulations are from the previous step of producing ‘O’ dsDNA and snapshot selection for ‘T’ dsDNA described above. As for ‘O’ dsDNA, over the 14,000 frames of the simulations, we selected the 8 following frames, ranging that we call O1 to O8, respectively: 6983, 7269, 7712, 8272, 9040, 10145, 10394, 11433. For UDG, the structure has been predicted by AlphaFold2 (AF2) with the isomer sequence from Uniprot ID P13051-2. The 56 first residues were discarded as they are in a disordered region with no confidence in its position. This structure is represented in Figure 5 with PyMOL^36^.

### Measurements of angles and distances in the base-flipped conformations

The opening angles for the flipped-out uracil bases in Table 1 are determined via the collective variable approach, as defined in the text. The method has been used for both the trinucleosome (‘T’) and control (‘C’) structures. The *d*_*U*−*groove*_ measure is calculated between the flipping out uracil and the nearest groove’s extremity. This distance spans between the N1 uracil atom and the nearest phosphate from the opposite strand. For an illustration, see Supplementary Figure 3 a), in which four positions P1, P2, P3, and P4 are defined representing the pseudo-dihedral points used to measure the uracil flipping angle around its backbone during the simulation.

### UDG flexibility

Flexibility was predicted using the MEDUSA webserver^38^, https://www.dsimb.inserm.fr/MEDUSA/. It predicts flexibility from the protein amino acids sequence in different modes. Each of these modes assign a probability of belonging to a predefined number of flexibility class per residue. The mode we have used is 3 class prediction; the results are summarized in Figure 5 a) and b). The structure in Figure 5 a) is represented with PyMOL^36^, Figure 5 b) is drawn with SSDraw^37^. The flexibility classes assigned according to MEDUSA are used as coloring factors; the explanation is given in the Figure caption.

### Selection of protein-DNA docking software

In order to determine which was the best-suited available software for UDG-dsDNA docking, we have tested different types of docking software. The complete list is given by: HDOCK^41^, HADDOCK^54^, pyDockDNA^43^, MDockPP^55^, RoseTTAFold2NA^56^ and AlphaFold3^57^. They were all evaluated on reproducing the interaction in the 1EMH crystal structure. From our tests, pyDockDNA performed the best on our use case as it discriminates between Uracil and Thymine, thus being the one selected to run the docking simulations. The summary of the comparison is found in Supplementary Figure S4.

### Evaluation of the docking structures

This protocol takes the 1EMH structure as a reference for the highest docking quality; it is the target which we wanted to reproduce by docking, and putting it more precisely, we measured the similarity with which the uracil enters the UDG compared to 1EMH. After aligning 1EMH UDG and the UDG of every docking conformation for each DNA strand with different base flipping angle and protocol, we measured the distance between the N3 atoms of the evaluated uracil (U1) and the uracil analog 2’-deoxypseudouridine of the reference uracil (U2). This *d*_*U*1−*U*2_ distance (in Å) is a direct quantitative evaluation of how much the docking conformation corresponds to the experimental structure. Figure 6 a) and b) both represent this evaluation performed on the trinucleosomic structure (‘T’ in a) and the oligomeric structure ‘O’ in b). The goal of this measure was to highlight which DNA strand features are impacting UDG action the most, thus the most successful docking conformation for each strand was taken and its quality was classified into ‘None’, ‘Low’, ‘Medium’ and ‘High’ quality according to the thresholds:

- ‘None’: *d*_*U*1−*U*2_ > 9 Å
- ‘Low’: 9 Å > *d*_*U*1−*U*2_ > 6 Å
- ‘Medium’: 6 Å > *d*_*U*1−*U*2_ > 3 Å
- ‘High’: *d*_*U*1−*U*2_ < 3 Å

## Data and code availability

All data and scripts of the article are hosted on this GIT repository: https://gitlab.in2p3.fr/cmsb-public/dyprosome.

## Acknowledgements

This work has been supported by the ANR grant “Dyprosome” (ANR-21-CE45-0032-02). We thank the Mesocenter of the University of Lille for the High Performance Computing resources and services they provided for the calculations performed during this project.

## Author contributions statement

R.B. conceived the study, V.S. performed the Molecular Dynamics simulations, N.R. performed the docking simulations of UDG on base-flipped structures from trinucleosome arrays and oligomeric DNA, G.B. provided advice for MD simulations and protein docking simulations; F.C. provided the discussion and MC simulations for the UDG search process on DNA described in the Supplementary Material and selected the trinucleosome linker structures from our earlier work. V.S., N.R., G.B., S.G., M.L., F.C. and R.B. analysed and discussed the results. All authors contributed to the writing of the manuscript.

## Supplementary Material

**ABSTRACT**

This Supplementary Material contains Supplementary Text and Figures to accompany the Main Text.

### An estimate of the efficiency of the uracil search process by UDG

About 200 to 500 C-to-U deaminations per cell per day are thought to occur naturally, both by exogenous and endogenous chemical oxidation events [S1]. We may consider that upon therapeutic irradiation such a number could increase even ∼100-fold, mostly by the action of free water-radicals. The average daily concentration of DNA-U in the typical cell nucleus (∅ ∼7 mm, V ∼1.4 pL) can therefore expected to be about 500 to 50,000/(6,02 ·10^23^) = 8·10^−22^ to 10^−20^ mol per 1.4 pL, that is 6 10−10 to 6 ·10−8 M.

UDG is found in the cell nucleus in concentrations that can vary between 0.1 to 10 ng/ml, according to the sensitivity range ELISA lab assays. Hence, a typical nucleus should contain a few 10^−16^ grams of UDG, with a molar weight of 25,7 kDa, that is order of 2,000 to 100,000 copies of the protein, also fluctuating during the cell cycle phases [S2]. Such a concentration seems nicely adjusted to match a ratio of roughly 1 UDG per uracil/day. However, if the absolute concentration could be about right, the main question is how UDG can identify the defective base among the 6 billion nucleotide pairs, that is the kinetics and the success rate of the repair process.

Experiments can track the kinetics of association of UDG with properly prepared DNA samples. For example, Tainer et al.S3] measured by stopped-flow electrophoresis the excision kinetics of UDG against small (19-nt), linear DNA fragments, either pristine, or including U:A and U:G mispairs. They stocked 8.5 pmol of DNA (that is, about 5 ·10^12^ copies of the oligonucleotide) and 0.02-0.04 ng of UDG (that is, about 5-10·10^8^ copies) in a reaction volume of 40 *μ*l. To compare with the in-vivo conditions, one may note that the DNA (besides being fragmented into tiny oligomers, and without histones) is diluted to about 2,000 times smaller than in the cell nucleus, whereas the UDG concentration is diluted to 2,500, so the relative proportions of UDG/DNA are roughly respected. However, the concentration of uracil is unrealistically high, since in the experiments there is one U every about 20 nt-pairs, although the DNA is broken and not continuous. On a proportional basis, this corresponds to about 1 UDG per 10,800 DNA-U defects, and is comparable to concentration ratios used in other similar experiments, see, e.g., [S3].

**Figure S1.**
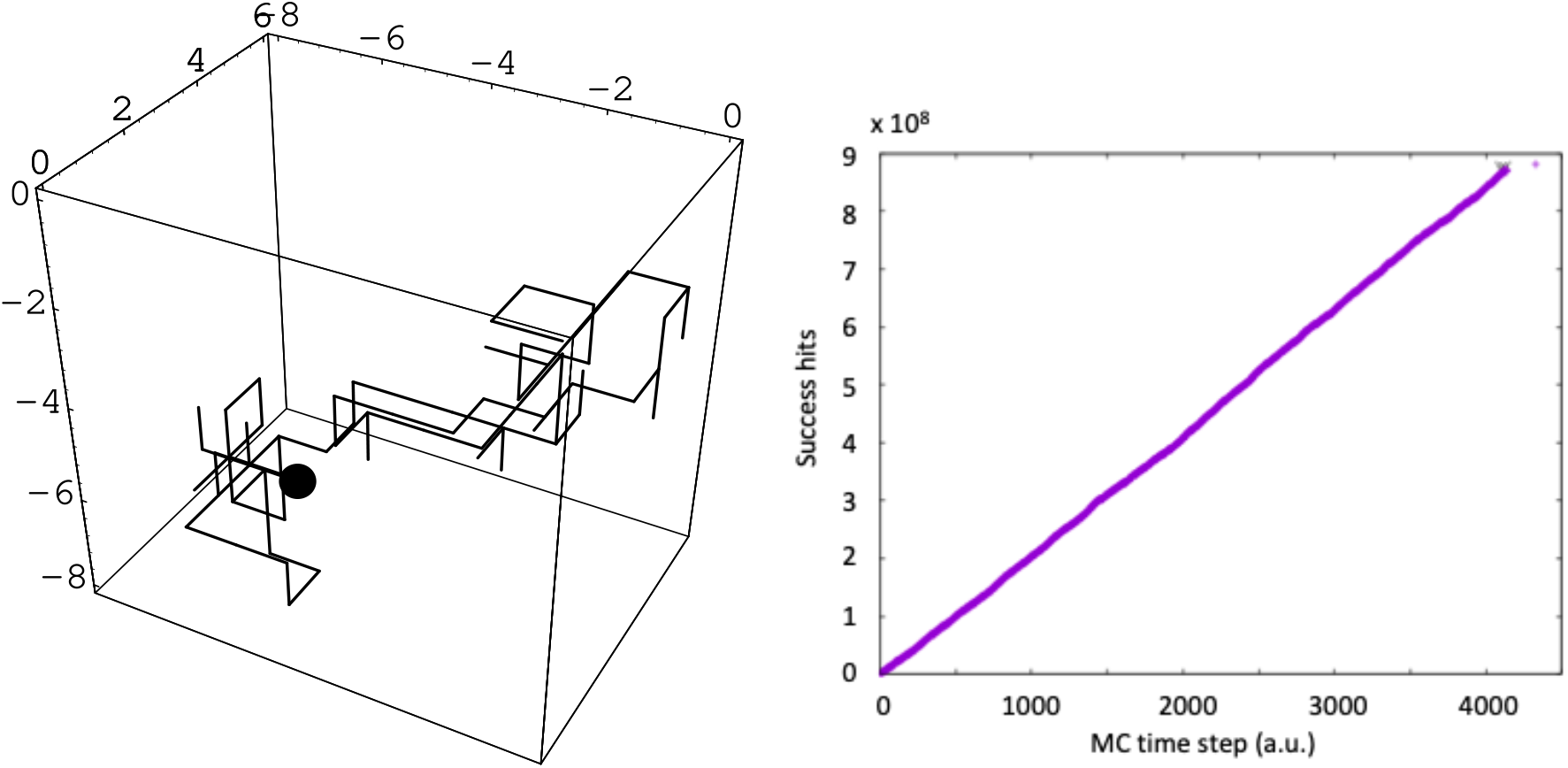
Figure 1 a) UDG performing a Brownian random walk; 1 b) Total number of success hits as a function of simulation time.

We ran a very simple Monte Carlo random search simulation, with the aim of estimating hitting times and search times of a single UDG along a fragment of DNA. We represented the search space as a dense collection of small cubic volumes of side 7 nm. Since the size of the globular UDG protein is roughly 5 nm, we imagine that each small cube is large enough to accommodate one copy of UDG and one fragment of DNA, in which case we count the couple as a successful identification of the uracil by UDG. Given the above concentration of 1 UDG per 10,800 DNA-U, it is enough to consider a space of (650)^3^ = 2.75 ·10^8^ cubes, including 10 800 randomly placed DNA at fixed positions, and one UDG performing a Brownian random walk, as shown in Fig. S1 a). At each MC time step the UDG jumps from one cube to another adjacent cube, also including stepping back to the starting one. Whenever UDG finds a DNA-U in a volume, a successful hit is counted. The slope of the linear plot in Fig. S1 b), displaying the total number of success hits as a function of the fictitious simulation time, gives a recursion time Δ t ∼25 000 Brownian steps, between two successive hits of UDG on two distinct DNA oligomers. The effective diffusion constant in water (with a kinetic viscosity *η* = 9.3 ·10^−4^ Pa s) of a globular protein with the size R = 2.5 nm of UDG, can be estimated by the Einstein relation

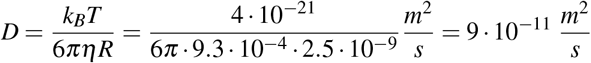

from which the elementary diffusion time per each step of 7 nm between two cubes, is estimated as *τ*≈ 0.54 *µs*. Hence, a Δ*t* ∼25000 *τ* ∼13.5 ms is the average time between each two successful hits on two distinct DNA oligomers, at the experimental concentrations.

By looking at the linear part of the kinetic plot reported in Tainer’s paper ([S3], Figure 1) a time of about ∼5 minutes is required to excise ∼180 *µ*mol U per mg of UDG, or 4,600 Uracil per UDG, that is about 65 ms for each event. The difference of about 50 ms between our calculated hitting time of 13,5 ms, and the excision time of 65 ms, appears to support the model put forward by the authors, according to which the UDG must firstly attach at a random location along the DNA oligomer, and then spends time to migrate along the length, “searching” for the defective nucleotide pair.

However, while such a picture seems to hold well for the experimental situation in which just 10-20 bases have to be searched in each fragment, it turns to fully unrealistic when translated to the in-vivo situation. Even neglecting the presence of densely packed chromatin into millions of nucleosomes, and taking the DNA as a free long, continuous polymer, at 10-to 100,000 uracil per 6 billion nt-pairs, each UDG should scan millions of nucleotide pairs according to this model: if 50 ms are needed to scan just about 20 nt-pairs in the experiment, finding one uracil by similarly scanning several million nt-pairs would require a time of days.

---

**Figure S2.**
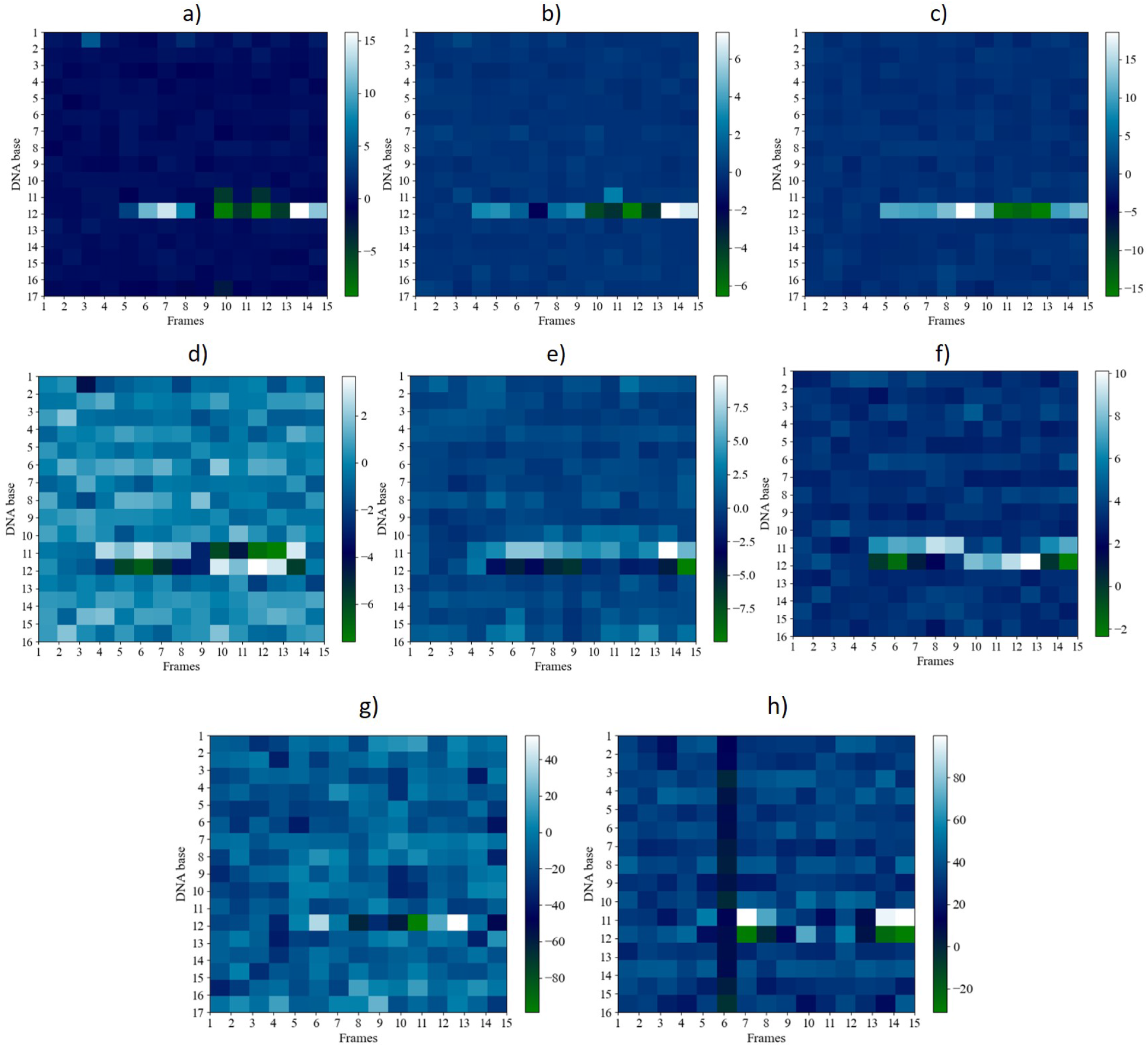
Intra-base pair parameters a) shear, b) stretch, c) stagger, g) propel, and inter-base pair parameters d) shift, e) slide, f) rise, and h) twist of uracil-mutated dsDNA for the CPDb scheme were calculated using the CURVES+ program. Here, the dark shades (blue to green) indicate negative values, and the light shades (cyan to white) indicate positive values of uracil-mutated dsDNA.

**Figure S3.**
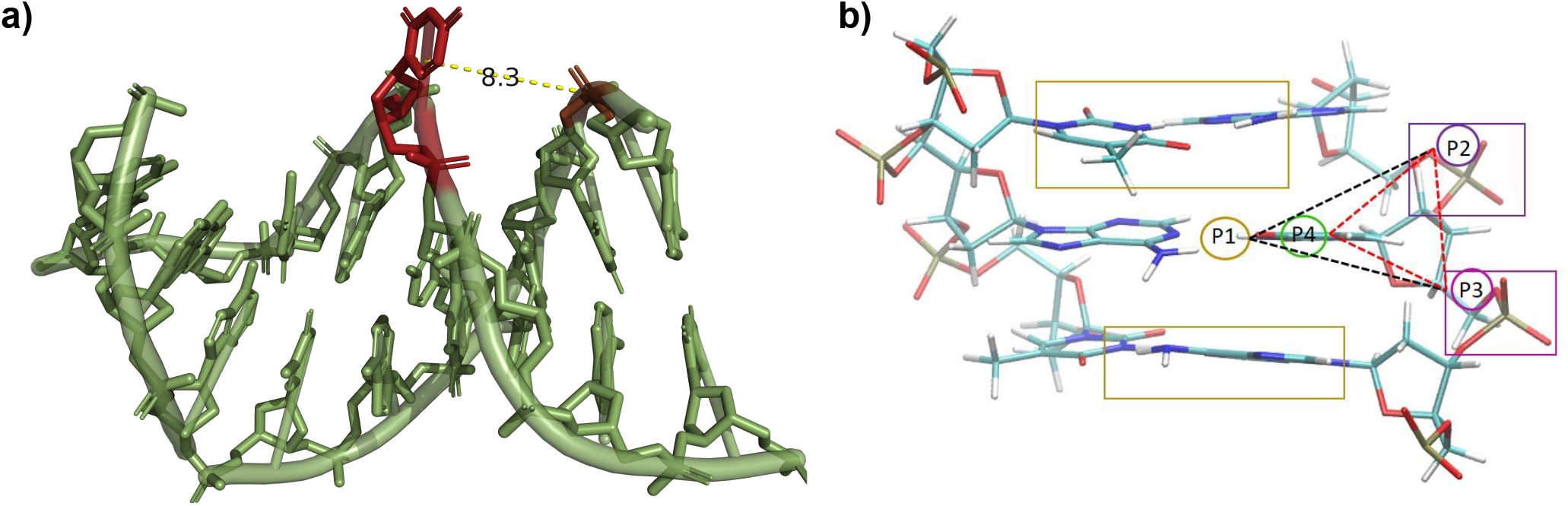
Illustration of the DNA feature measurements listed in Table 1; a) Distance measured (Å) in structure T10 (green; two Pymol representations - sticks and cartoon (with transparency)) between the position N1 of the flipped out uracil (red) and the P atom of the closest phosphate group (brown) of the opposite strand’s backbone. b) Here, point P1 is the center mass of two flanking base pairs (brown color rectangular box: residues 11, 13, 22, and 24), point P2 (purple rectangular box) and point P3 (magenta rectangular box) are defined by the flanking phosphate group, and point P4 is defined by the target base, i.e. uracil for the flipping process. The dashed lines define the pseudo-dihedral angle formed by four points P1, P2, P3, and P4 measured between the planes passing through P1, P2, P3 (black) and P2, P3, P4 (red).

**Figure S4.**
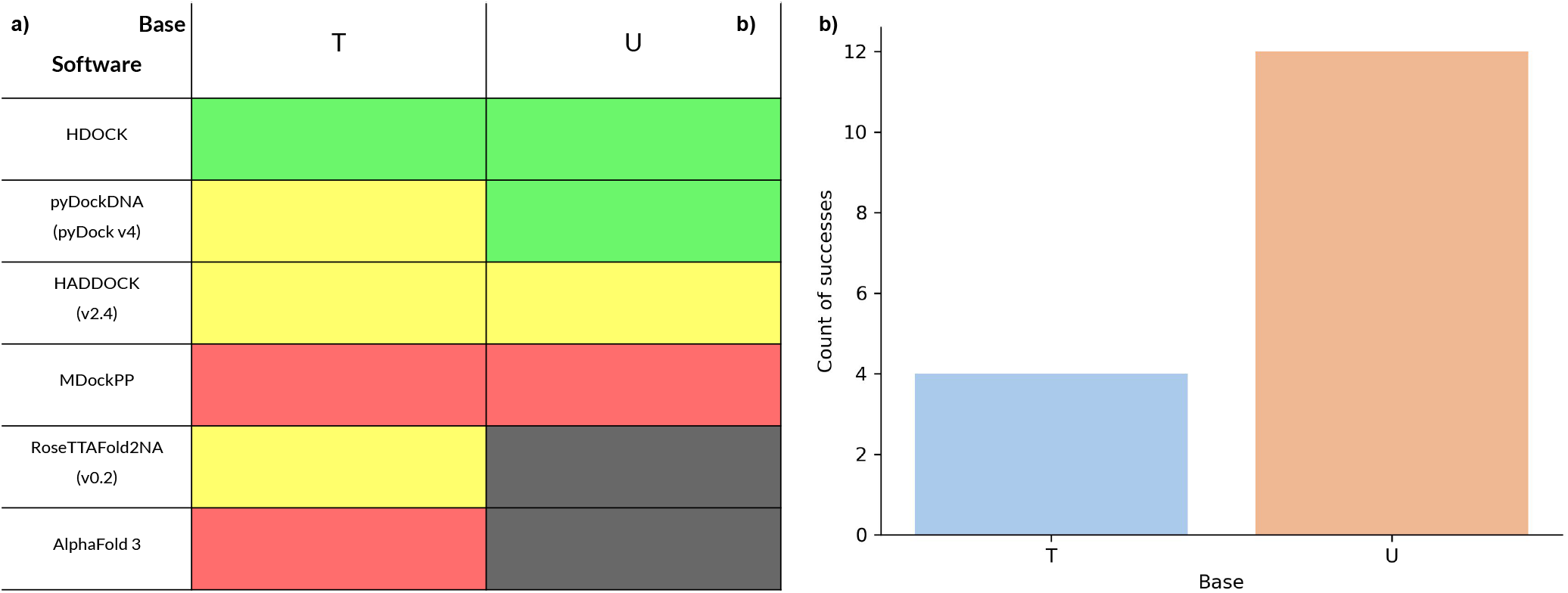
a) Table reporting the success of reproducing the 1EMH structure by docking. Inputs given to the software are the separated chains from the complex (for the first four programs): the dsDNA, either with the uracil analog replaced with a thymine or a uracil, and the UDG structure from the crystal structure. Sequences were provided to the last two software packages that do not take structures as input. The color code is the following: <u>green</u> for complete success, <u>yellow</u> for partial success (e.g., the protein-DNA interface is conserved but the substrate residue is not predicted inside the catalytic pocket), <u>red</u> for failure and <u>black</u> for software-inherent inability. See main text for the definition of success. b) pyDockDNA success counts for the DNA substrate containing a uracil compared to when it contains a thymine, justifying the ‘partial success’ for pyDockDNA with a thymine in a).

**Figure S5.**
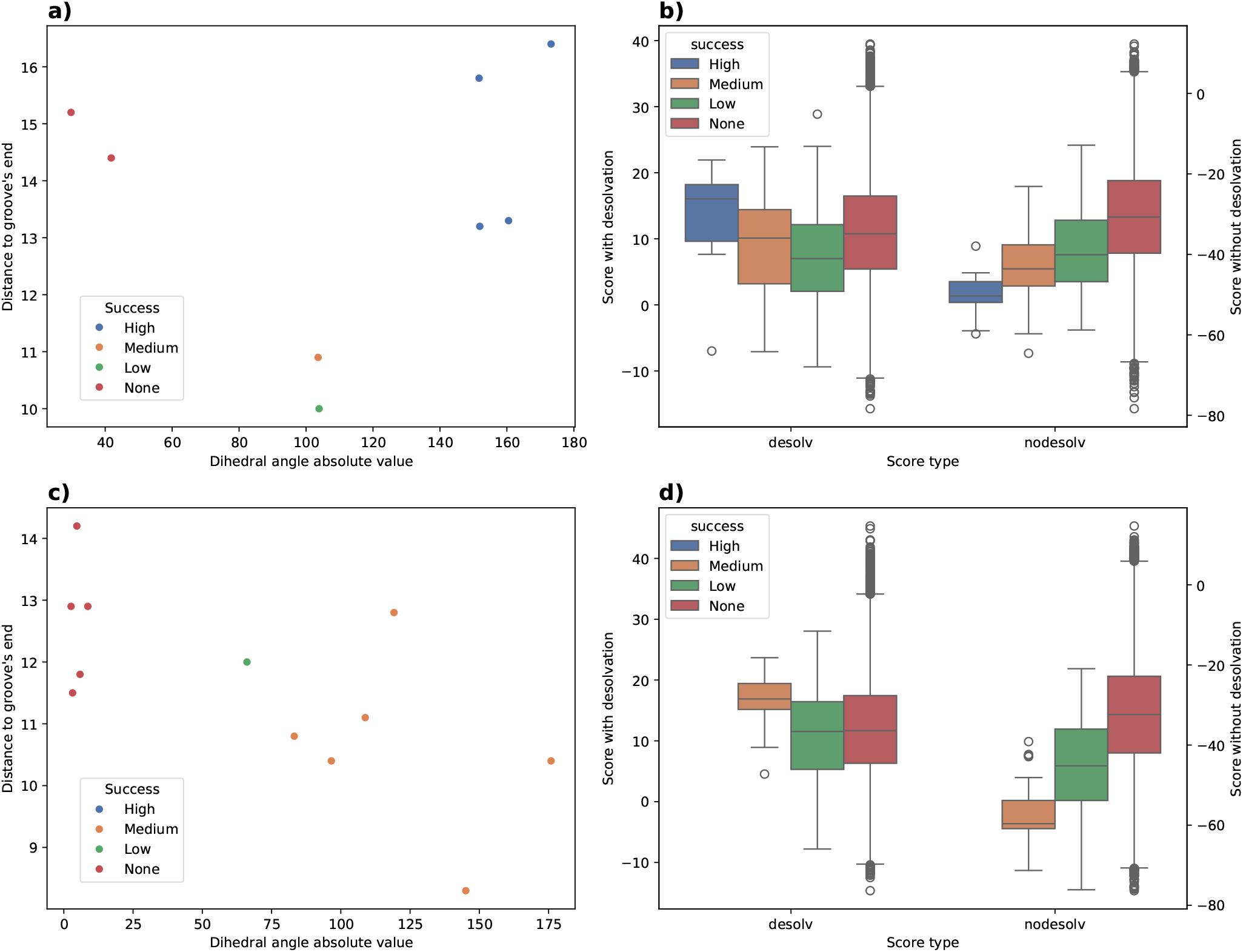
Supplementary information on docking quality. Figures a) & b) refer to oligomeric - ‘O’-dsDNA, Figures c) & d) are for trinucleosomic - ‘T’ - dsDNA. For the two dsDNA types, two different types of plots are displayed. The first type, in a) and c), shows scatter plots for docking success as a function of the base flipping angle measured by the dihedral angle and the groove width, represented by the distance from the uracil to the nearest groove’s end; the second type, in b) and d), show boxplots of the distribution of pyDockDNA predicted score (with or without desolvation) depending on the finally evaluated docking success.

**Figure S6.**
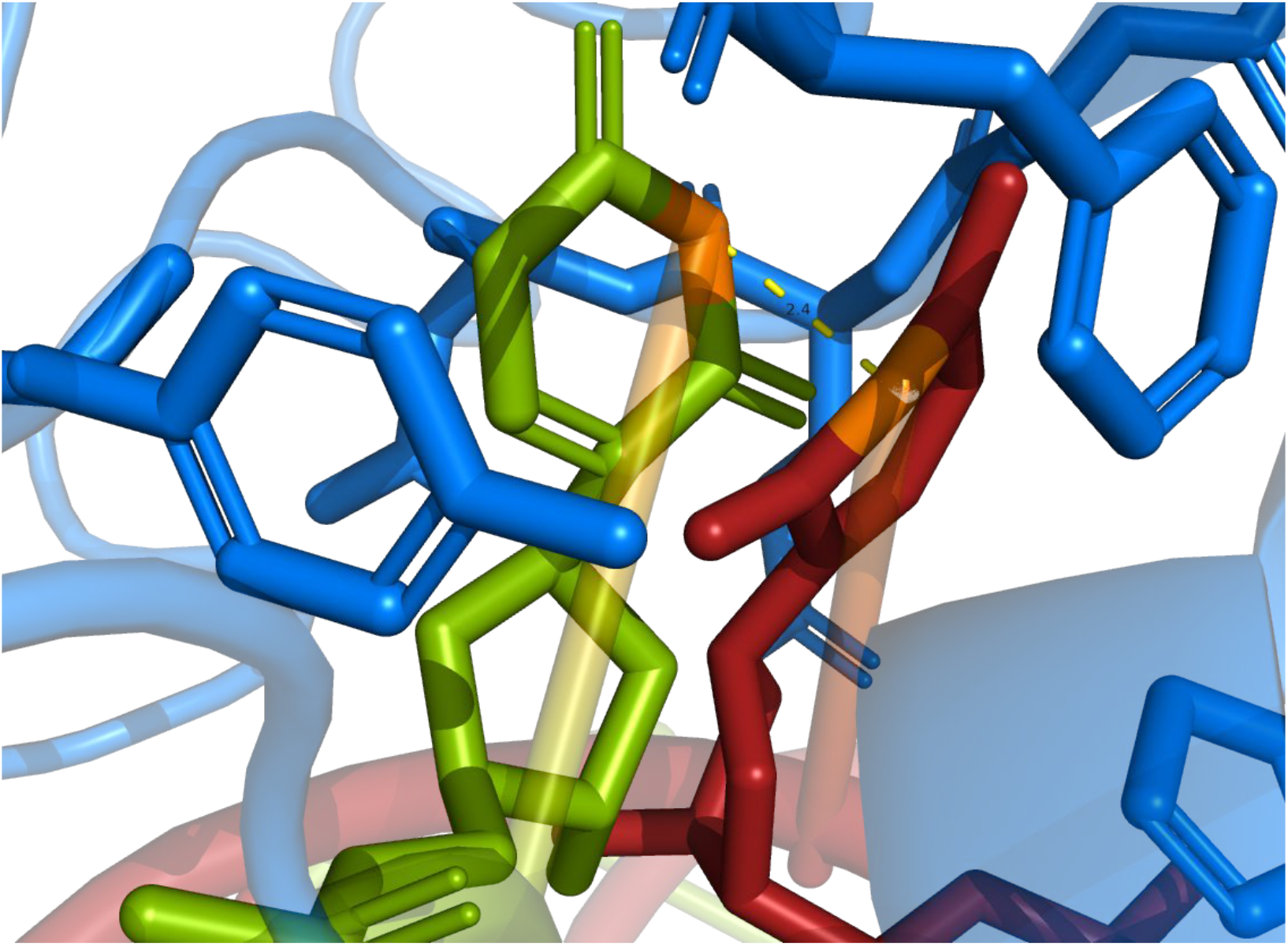
Representation of the evaluation for ‘O6’ best docking conformation by distance measurements. The figure is visualized with PyMOL and respects the following color code and representation: the dsDNA is represented in a cartoon style with their uracil (or uracil analog) as sticks. 1EMH dsDNA is in green while the ‘O6’ dsDNA is in red. UDG is in blue and cartoon except for its catalytic pocket being represented as sticks. Transparency is at 50% applied on the cartoon representation. Atoms located at the endpoint of the measurement lengths are colored in orange. The distance between ‘O6’ uracil and 1EMH uracil analog 2’-deoxypseudouridine is computed between their N3 atoms.

**Figure S7.**
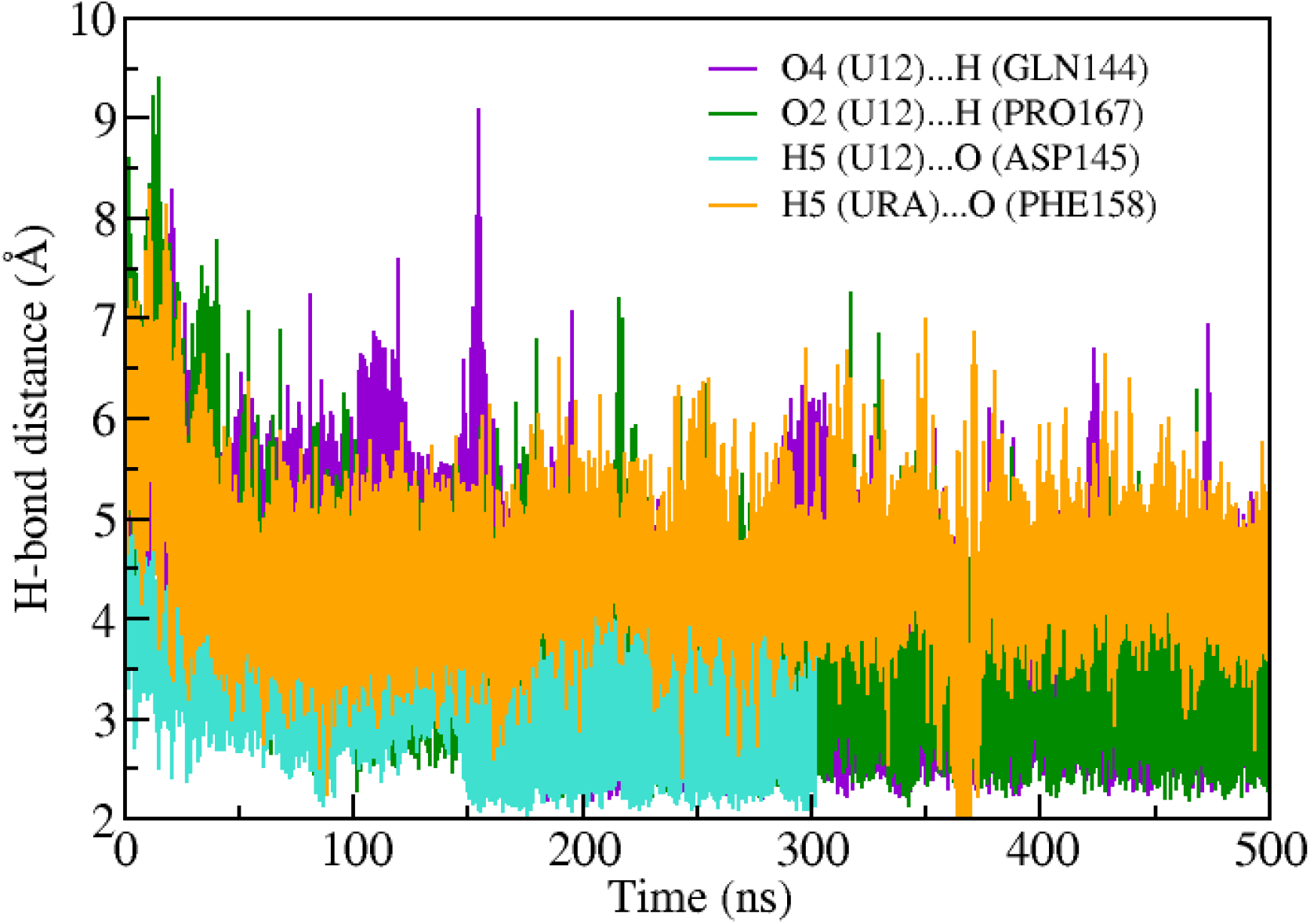
Hydrogen bond distance between the uracil and the catalytic sites of UDG enzyme

